# Genome-wide association study meta-analysis of the Alcohol Use Disorder Identification Test (AUDIT) in two population-based cohorts (N=141,932)

**DOI:** 10.1101/275917

**Authors:** Sandra Sanchez-Roige, Abraham A. Palmer, Pierre Fontanillas, Sarah L. Elson, The 23andMe Research Team, Substance Use Disorder Working Group of the Psychiatric Genomics Consortium, Mark J. Adams, David M. Howard, Howard J. Edenberg, Gail Davies, Richard C. Crist, Ian J. Deary, Andrew M. McIntosh, Toni-Kim Clarke

## Abstract

Alcohol use disorders (**AUD**) are common conditions that have enormous social and economic consequences. We obtained quantitative measures using the Alcohol Use Disorder Identification Test (**AUDIT**) from two population-based cohorts of European ancestry: UK Biobank (**UKB**; N=121,604) and 23andMe (N=20,328) and performed a genome-wide association study (GWAS) meta-analysis. We also performed GWAS for AUDIT items 1-3, which focus on consumption (**AUDIT-C**), and for items 4-10, which focus on the problematic consequences of drinking (**AUDIT-P**). The GWAS meta-analysis of AUDIT total score identified 10 associated risk loci. Novel associations localized to genes including *JCAD* and *SLC39A13;* we also replicated previously identified signals in the genes *ADH1B, ADH1C, KLB*, and *GCKR*. The dimensions of AUDIT showed positive genetic correlations with alcohol consumption (r_g_=0.76-0.92) and Diagnostic and Statistical Manual of Mental Disorders (DSM-IV) alcohol dependence (r_g_=0.33-0.63). AUDIT-P and AUDIT-C showed significantly different patterns of association across a number of traits, including psychiatric disorders. AUDIT-P was positively genetically correlated with schizophrenia (r_g_=0.22, p=3.0×10^−10^), major depressive disorder (r_g_=0.26, p=5.6×10^−3^), and attention-deficit/hyperactivity disorder (ADHD; r_g_=0.23, p=1.1×10^−5^), whereas AUDIT-C was negatively genetically correlated with major depressive disorder (r_g_=−0.24, p=3.7×10^−3^) and ADHD (rg=−0.10, p=1.8×10^−2^). We also used the AUDIT data in the UKB to identify thresholds for dichotomizing AUDIT total score that optimize genetic correlations with DSM-IV alcohol dependence. Coding individuals with AUDIT total score of ≤4 as controls and ≥12 as cases produced a high genetic correlation with DSM-IV alcohol dependence (r_g_=0.82, p=3.2×10^−6^) while retaining most subjects. We conclude that AUDIT scores ascertained in population-based cohorts can be used to explore the genetic basis of both alcohol consumption and AUD.

## Introduction

Alcohol use disorders (**AUD**) are modestly heritable, with twin-studies demonstrating that approximately 50% of the phenotypic variance is attributed to genetic factors (1, 2). To date, genetic studies of AUD have identified genes that influence pharmacokinetic (e.g. *ADH1B, ADH1C*, ALDH2)(3–8), but not pharmacodynamic factors. The difficulty of assembling large, carefully diagnosed cohorts of AUD has stimulated additional studies of non-clinical phenotypes, such as alcohol consumption, in populations not ascertained for alcohol dependence. This approach has allowed for the relatively rapid collection of much larger sample sizes (e.g. >100,000s individuals) and has identified numerous loci associated with both pharmacokinetic and pharmacodynamic factors that influence alcohol consumption, including *ADH1B/ADH1C/ADH5* (9–11), *KLB* (encoding β-klotho)(9, 11, 12) and *GCKR*, encoding the glucokinase regulatory protein (9, 11). However, the genetic overlap between alcohol consumption (units per week) and DSM-IV diagnosed alcohol dependence is moderate (r_g_ = 0.38)(13), reinforcing the notion that alcohol consumption cannot be used as a surrogate for alcohol dependence or AUD.

The Alcohol Use Disorders Identification Test (**AUDIT**) is a screening tool designed to identify hazardous alcohol use in the past year (14). The test consists of 10 items across 3 dimensions pertaining to alcohol consumption (items 1-3, often termed **AUDIT-C**), dependence symptoms (items 4-6), and harmful alcohol use (items 7-10) (collectively **AUDIT-P**). When the AUDIT was developed, a total score of 8 or higher was proposed to be indicative of harmful alcohol use (14) and a score of 20 or higher consistent with a diagnosis of alcohol dependence (15); however, there is no clear consensus and subsequent studies have suggested that additional factors including sex and cultural and social contexts should be considered when deriving thresholds for alcohol dependence (reviewed in **Supplementary Table 1**).

A recent population-based GWAS of AUDIT in 20,328 research participants from the genetics company 23andMe, Inc., identified a locus near the gene *ADH1C* (rs141973904; p = 4.4 × 10^−7^)(10) nominally associated with AUDIT total score. AUDIT scores among 23andMe research participants were low and predominantly driven by alcohol consumption (**AUDIT-C**). The genetic correlation between AUDIT total score from 23andMe and alcohol consumption was much stronger (r_g_ = 0.89, p = 9.01 × 10^−10^) than the genetic correlation between AUDIT total score and alcohol dependence (r_g_ = 0.08; p = 0.65)(13).

In this study, we performed a GWAS meta-analysis using the UK Biobank (**UKB**; N = 121,604) and the previously published 23andMe cohort (N = 20,328)(10), yielding the largest GWAS meta-analysis of AUDIT total score to date (N = 141,932). Using only the UKB cohort, we also sought to determine whether the alcohol consumption component of the AUDIT had a genetic architecture distinct from the dependence and harmful use components by performing GWASs of consumption “AUDIT-C” (items 1-3) and problems “AUDIT-P” (items 4-10). Linkage Disequilibrium Score regression (**LDSC**)(16) was used to calculate genetic correlations between AUDIT measures and other substance use, psychiatric, and behavioral traits. We also calculated genetic correlations with obesity and blood lipid traits, as these have previously been shown to associate with alcohol consumption (9, 10). We hypothesized that AUDIT-P would correlate more strongly with measures of hazardous substance use, including alcohol dependence, and other psychiatric conditions. Finally, in order to determine the thresholds for dichotomizing AUDIT total score that would most closely approximate alcohol dependence, we converted continuous AUDIT total score into cases and controls using different thresholds, performed GWAS on each, and calculated the genetic correlation with DSM-IV alcohol dependence (13).

## Materials and Methods

### UK Biobank sample

The UK Biobank (**UKB**) is a population-based sample of 502,629 individuals who were recruited from 22 assessment centers across the United Kingdom from 2006-2010 (17). 157,366 individuals completed a mental health questionnaire as part of an online follow-up over a one-year period in 2017. The Alcohol Use Disorder Identification Test (**AUDIT**)(14) was administered to assess alcohol use over the past year, using gating logic (see **Supplementary Figure 1**). After performing quality control to remove participants with missing data, and keeping only white British unrelated individuals, 121,604 individuals with AUDIT total scores were available. AUDIT total score was created by taking the sum of items 1-10 for all participants, including those who endorsed currently never drinking alcohol (as they could still endorse past alcohol harm on items 9 and 10). We also created AUDIT subdomain scores by aggregating the scores from items 1-3, which include the information pertaining to alcohol consumption (**AUDIT-C**, N = 121,604), and from items 4-10, which indexes the information pertaining to alcohol problems (**AUDIT-P**, N = 121,604). These traits were log10 transformed to approximate a normal distribution (**Supplementary Figure 2**).

### Genotyping, quality control and imputation

Genotype imputation was performed on 487,409 individuals by the UKB team using IMPUTE4 (18) and the Haplotype Reference Consortium reference panel. After quality control, 16,213,998 SNPs remained for GWAS analyses. Additional details on genotyping and quality control are shown in the **Supplementary Material**.

### Discovery GWASs using UKB

GWAS analyses were performed using BGENIE v1.1 (18) with AUDIT scores (AUDIT total score, AUDIT-C, and AUDIT-P, tested independently) as the outcome variable and age, sex, genotyping array and the first 20 principal components derived from genotype data as covariates. See the **Supplementary Material** for extended details. In order to identify independently-associated variants (“index variants”), clump-based pruning was applied in FUMA using an r^2^ of 0.1 and a 1 Mb sliding window using the UKB White British sample as the LD reference panel. A 1 Mb window was used due to the regions of extended linkage disequilibrium on chromosomes 4q23 and 17q21.31, which were associated with AUDIT scores in this study.

In addition, we performed a series of 18 case-control GWAS analyses of AUDIT total score using different thresholds (cases: ≥8, 10, 12, 15, 18, 20 vs controls: ≤2, 3, 4). The sample size at each threshold is shown in **Supplementary Table 2**. The results of these analyses were used to determine which thresholds would produce the highest genetic correlation estimates with DSM-IV defined alcohol dependence (13).

### SNP-Heritability analyses

The SNP-heritability of UKB AUDIT scores (total, AUDIT-C, AUDIT-P) was calculated using a genomic restricted maximum likelihood (**GREML**) method implemented in Genetic Complex Trait Analysis (**GCTA**)(19) on a subset of 117,072 unrelated individuals using a relatedness cutoff of 0.05 and controlling for age and sex. GREML analyses were run using genotyped SNPs with a MAF greater than 0.01 to construct the GRM.

### GWAS meta-analysis of AUDIT total score using the UKB and 23andMe cohorts

Because the genetic correlation of AUDIT total score between the UKB and 23andMe cohorts was high (r_g_ = 0.77, SE = 0.12, p = 7.15 × 10^−11^), we performed a sample-size based meta-analysis of AUDIT total score from the UKB and 23andMe cohorts using METAL (version 2011-03-25)(20). This meta-analysis comprises a total of 141,932 research participants of European ancestry and 9,519,872 genetic variants that passed quality control. We used clump-based pruning (see ‘Discovery GWAS’) to identify independently-associated variants. For each GWAS signal we defined a set of credible variants using a Bayesian refinement method developed by Maller et al. (21). These credible sets are considered to have a 99% probability of containing the ‘causal’ variant at each locus. Credible set analyses were performed in R (https://github.com/hailianghuang/FM-summary) for each of the index variants associated with AUDIT total score in the GWAS meta-analysis using SNPs within 1 Mb with an r^2^ >0.4 to the index variant. All downstream genetic analyses of AUDIT total score were performed using the GWAS meta-analysis summary statistics. The 23andMe AUDIT GWAS has previously been published (10) and 30,441 participants from the UKB cohort were included in a previous GWAS of alcohol consumption (9).The GWAS of alcohol consumption in UKB involved individuals genotyped during the first UKB data release (9). The individuals from UKB in the current study are those who completed the mental health questionnaire in 2017 and include those genotyped in both first and second UKB data release; the sample overlap between these two datasets is 30,441 individuals.

### Functional mapping and annotation of GWAS meta-analysis

We used FUMA v1.2.8 (22) to study the functional consequences of the index SNPs, and of the SNPs contained in each credible set, which included ANNOVAR categories, Combined Annotation Dependent Depletion (**CADD**) scores, RegulomeDB scores, eQTLs, and chromatin states. We also studied the regulatory consequences of the index SNPs using the Variant Effect Predictor (**VEP**; Ensembl GRCh37).

### Gene-set and pathway analyses

We performed MAGMA (22) competitive gene-set and pathway analyses using the summary statistics from the GWAS meta-analysis of AUDIT total score and the AUDIT-C and AUDIT-P subsets. SNPs were mapped to 18,546 protein-coding genes from Ensembl build 85. Gene-sets were obtained from Msigdb v5.2 (“Curated gene sets”, “GO terms”).

### Gene-based association using transcriptomic data with S-PrediXcan

We used S-PrediXcan (23) to predict gene expression levels in 10 brain tissues, and to test whether the predicted gene expression correlates with AUDIT scores. We used pre-computed tissue weights from the Genotype-Tissue Expression (**GTEx** v7) project database (https://www.gtexportal.org/) as the reference transcriptome dataset. Further details are provided in the **Supplementary Material**.

### Genetic correlation analysis

We used LD Score regression (**LDSC**) to identify genetic correlations between traits (24). This method was used to calculate genetic correlations (r_g_) between AUDIT total score, AUDIT-C, and AUDIT-P and 39 other traits and diseases (see **Supplementary Tables 3, 4** and **5**). We did not constrain the intercepts in our analysis, as we could not quantify the exact amount of sample overlap between cohorts. We used False Discovery Rate (**FDR**) to correct for multiple testing (25). We also used LDSC to examine genetic correlations between various dichotomized versions of AUDIT and DSM-IV defined alcohol dependence (13). To test for significant differences between the genetic correlations, z-score statistics were calculated (see **Supplementary Table 6**).

## Results

### UKB sample demographics and characteristics

In the UKB cohort, there were 121,604 individuals with AUDIT scores available for GWAS analysis (**Supplementary Table 7**). The UKB sample was 56.2% female (N = 68,389) and the mean age was 56.1 years (S.D. = 7.7). The mean AUDIT total score was 5.0 (S.D. = 4.18, range = 0-40); a histogram showing the distribution of the scores is shown in **Supplementary Figure 2**. Over the prior year, 91.9% of the participants reported drinking 1 or 2 drinks on a single day. Over the prior year, 6.3% of the participants reported they were not able to stop drinking once they started, and 10.7% felt guilt or remorse after drinking alcohol (**Supplementary Table 7**). Males had significantly higher AUDIT total mean scores than females (6.09 ± 4.45 vs. 4.15 ± 3.72, respectively; β = 0.47, p < 2 × 10^−6^; **Supplementary Figure 3**). In addition, age was negatively correlated with AUDIT scores (β = −0.02, p < 2 × 10^−6^; **Supplementary Table 8**). Therefore, both sex and age were used as covariates in the GWAS analyses. The mean AUDIT-C score was 4.24 (S.D. = 2.83) and the mean AUDIT-P score was 0.75 (S.D. = 2.0). As expected, there was a moderate positive phenotypic correlation between AUDIT-C and AUDIT-P (r = 0.478, 95% C.I. = 0.473-0.481, p < 2 × 10^−16^; **Supplementary Table 8**).

### SNP-heritability in UKB

We estimated the SNP-heritability of AUDIT total score to be 12% (**GCTA**: ± 0.48%, p = 4.6 × 10^−273^; **LDSC**: 8.6% ± 0.50%), which is similar to the estimate from Sanchez-Roige et al. (10). The SNP-heritability for AUDIT-C was 11% (**GCTA**: ±0.47%, p = 1.5 × 10^−211^; **LDSC**: 8.4% ±0.55%), and 9% for AUDIT-P (**GCTA**: ±0.46%, p = 2.0 × 10^−178^; **LDSC**: 5.9% ±0.48%).

### GWAS of AUDIT scores in UKB

The significant results (p < 5 × 10^−8^) of the GWAS of AUDIT total score in the UKB cohort are shown in **Supplementary Table 9**; this analysis revealed 12 independent GWAS signals located in 8 loci. The UKB GWAS of AUDIT-C and AUDIT-P subsets are summarized in **Supplementary Tables 10** and **11** and **Supplementary Figure 4**. Seven of these 12 independent GWAS signals also significantly associated with AUDIT-C; the same index variants were identified in the two analyses. An additional GWAS signal was also identified close to *FNBP4*. For AUDIT-P, 5 independent GWAS signals were significantly associated and these loci were also associated with the total AUDIT and AUDIT-C. The rs1229984 SNP in *ADH1B* was not available for meta-analysis in the 23andMe sample and was not in Hardy Weinberg equilibrium (**HWE**) in the UKB sample used in the present study (p = 3.2 × 10^−16^); however, in the total UKB White British sample there was no significant deviation from HWE (p = 0.13). The association between rs1229984 and AUDIT scores are presented in **Supplementary Tables 9, 10** and **11**. rs1229984 was strongly associated with all AUDIT scores in the UKB (β = 0.04-0.06, p < 1.0 × 10^−45^), but this SNP was not included for clump-based pruning and downstream analyses. As such, we performed a conditional analysis of the SNPs on 4q23 and 4q24 in the UKB sample to determine whether any of these associations were significant after controlling for rs1229984 genotype. While rs13107325 on 4q24 remained significantly associated with AUDIT total score after controlling for rs1229984 genotype, the association between rs146788033, rs11733695 and rs3114045 and AUDIT total score became non-significant, suggesting that these loci are tagging the strong rs1229984 signal in this region.

### GWAS meta-analysis of AUDIT total score

The GWAS meta-analysis of the UKB and 23andMe samples found 15 independent GWAS signals (**Supplementary Table 12**) associated with AUDIT total score spanning 10 genomic loci (**Table 1**). Figure 1 shows the Manhattan and QQ plots of the GWAS meta-analysis of AUDIT total score and **Supplementary Figures 5-14** show the regional association plots for the independent signals. The inflation factor of the meta-analysis GWAS was λ_GC_ = 1.22 with an LDSC intercept of 1.008 (SE = 0.007), suggesting that the majority of the inflation is due to polygenicity. The 15 independent SNPs show 100% sign concordance for association with AUDIT total score across UKB and 23andMe (**Table 1**); 11 of these SNPs were nominally associated with AUDIT total score in 23andMe (p ≤ 0.05), and all index SNPs were associated with AUDIT total score in UKB (p < 1.8 × 10^−6^).

**Figure 1.**
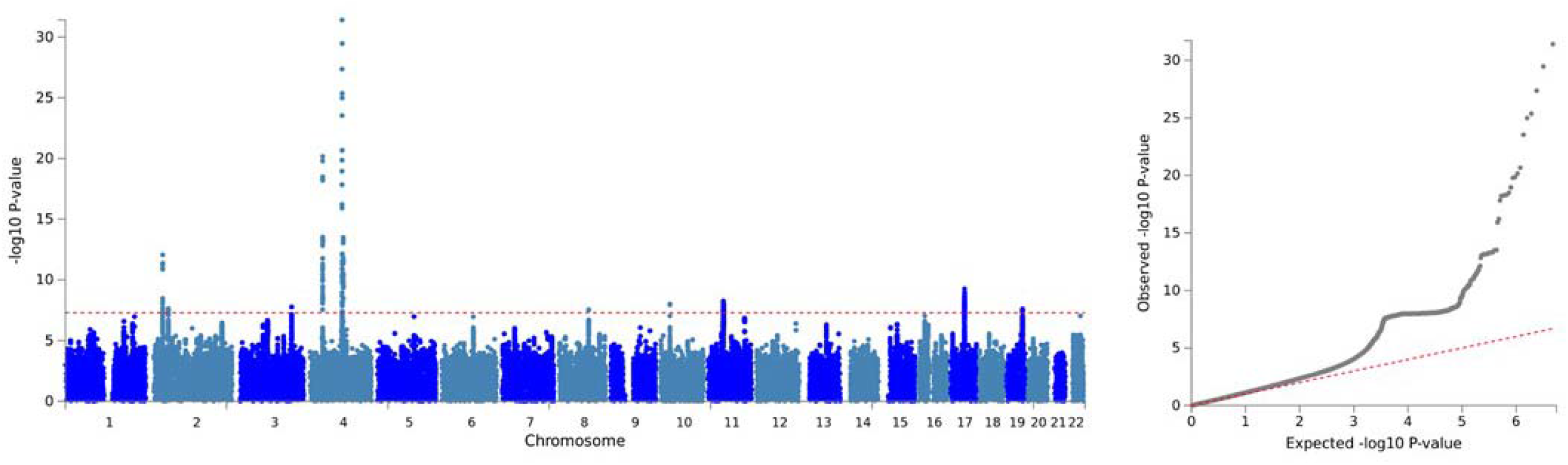
Manhattan and QQ plots for the SNP-based GWAS meta-analysis of AUDIT total score (N = 141,932)

The top hit for the GWAS meta-analysis of AUDIT total score was a variant (rs11733695) located downstream (879 base pairs) from *ADH6* (p = 3.4 × 10^−30^), which is a member of the alcohol dehydrogenase gene family. rs11733695 is in low LD (r^2^ = 0.17) with the functional SNP in *ADH1B*, rs1229984, which is known to alter alcohol metabolism (26). In addition, two other regions in 4q23 were associated with AUDIT total score in the meta-analysis: one of the index SNPs was located in the *ADH1C* gene; however, conditional analysis of this region in UKB alone suggests that these multiple hits may in fact be tagging the rs1229984 signal. This region has been previously associated with alcohol consumption, AUD, and AUDIT scores (6, 7, 9, 10)

We also replicated the association between *KLB* (**Supplementary Table 12**), on chromosome 4q14, and alcohol consumption (9, 11, 12); the index SNP rs11940694, which is located in the intron of *KLB*, was associated with AUDIT total score in the present study. Clump based pruning identified rs11940694 and rs4975012 as independent hits in the *KLB* region. Credible set analysis suggests that rs2046330 is the index SNP in the region represented by rs4975012 (**Supplementary Table 13**). AUDIT total score was also associated with SNPs that localized to *GCKR* on chromosome 2p23.3, which has been previously associated with alcohol consumption (9, 11). Five SNPs comprised the credible set at the *GCKR* locus, which also spans the *SNX17* gene, including the missense variant (rs1260326) in *GCKR* that was identified as the index SNP.

We identified GWAS signals in several regions that have not been previously implicated in the genetics of AUD, including 2p21, 17q21, 3q25, 8q22, 10p11, 11p11 and 19q13. The index SNP rs13135092 in the 4q24 region is located in an intron of *SLC39A8;* the remainder of the credible set for this locus is located in a non-coding pseudogene, *RN7SL728P. SLC39A8* is highly pleiotropic (27) but it is a novel association in relation to alcohol. A region of association on 2p21 contains 17 SNPs that are localized to the non-coding RNA, *LINC01833*. A novel region of association was also detected on chromosome 10p11.23; this region contains 9 credible SNPs that localize to the *JCAD* (junctional cadherin 5 associated) gene. *JCAD* encodes an endothelial cell junction protein, and has previously been associated with coronary heart disease (28).

The remaining novel associations on 3q25, 8q, 11p11, 17q21 and 19q13.3 were more complex. The index variants on chromosome 8q22.1 were not localized to any genes, and it is unclear from the credible set analyses what the causal variants may be at these loci. The credible SNP sets for the 3q25.33, 11p11.2, 17q21.31 and 19q13.3 regions contained over 50 SNPs each, which spanned several genes. For example, the index SNP on chromosome 17q21.31 was an intronic SNP in *MAPT*, which encodes the tau protein and has been robustly associated with Parkinson’s disease (30, 31) (**Supplementary Table 14**) and other neurodegenerative tauopathies (32), and more recently with neuroticism (33). However, we note that the region of association on chromosome 17q21.31 spans the corticotrophin receptor gene (*CRHR1*), which has been associated with alcohol use in animals and humans (34). Thus, due to the extended complex LD in this region, we are unable to determine the likely causal variant. Similarly, the index SNP (rs2293576) at chromosome 11p11.2 is a synonymous SNP of the zinc transporter gene *SLC39A13*; however, this region includes 54 associated SNPs, which map to five additional genes. Lastly, the index SNP on 19q13.3 is a synonymous variant in *FUT2. FUT2* encodes galactoside 2-alpha-L-fucosyltransferase 2, which controls the expression of ABO blood group antigens. However, 70 SNPs form this credible set and they span 4 genes in total.

We used FUMA to functionally annotate all 1,290 SNPs in the credible sets (see **Supplementary Table 13**). The majority of the SNPs were intronic (76.9%; N = 993) or intergenic (11.6%; N = 149), while only 26 SNPs (2.0%) were exonic. Furthermore, 38 SNPs showed CADD scores >12.37, which is the suggested threshold to be considered deleterious (35). The exonic SNPs (rs601338, rs17651549, rs13107325) of *FUT2, MAPT* and *SLC39A8*, respectively, had the highest CADD scores (>34), suggesting potential deleterious protein effects. 164 SNPs had RegulomeDB scores of 1a-1f, showing evidence of potential regulatory effects. 90.1% of the SNPs were in open chromatin regions (minimum chromatin state 1-7).

### Gene-based and pathway analyses

We used MAGMA (22) to perform a gene-based association analysis; which identified 40 genes that were significantly associated with AUDIT total score (p < 2.70 × 10^−6^; **Supplementary Table 15, Supplementary Figure 15**). As expected, the majority of these genes were in the 10 GWAS loci (i.e. *KLB, GCKR); DRD2* and *CRHR1* were also among the top hits. In addition, the analysis revealed a strong burden signal in *CADM2* (p = 1.64 × 10^−9^), where the index variant in GWAS meta-analysis did not reach genome-wide significance. We did not identify any canonical pathways that were significantly associated with AUDIT (**Supplementary Table 16**).

Gene-based (**MAGMA**) analyses for the AUDIT-C and AUDIT-P subsets (**Supplementary Figures 16** and **17**, respectively) revealed evidence of overlap (**Supplementary Figure 18, Supplementary Table 17**). Two genes (*KLB, CADM2*) were associated with all 3 AUDIT traits (AUDIT total score, AUDIT-C, and AUDIT-P). There was considerable overlap between AUDIT total score and AUDIT-C, with 21 overlapping genes associated at the gene-based level. Only one gene, *DRD2*, was associated with both AUDIT total score and AUDIT-P.

### S-PrediXcan

S-PrediXcan identified a positive correlation (p < 1.07 × 10^−6^) between AUDIT total score and the predicted expression of 26 genes across multiple brain tissues (full results are presented in **Supplementary Table 18**), including *MAPT* (cerebellum), *FUT2* (caudate and nucleus accumbens), and *CRHR1-IT1* (putamen, cerebellum, hippocampus, anterior cingulate cortex). SNPs in the region of *MAPT* and *FUT2* were associated with AUDIT total score in the GWAS. *MAPT* (cerebellum), *FUT2* (nucleus accumbens) and *CRHR1-IT1* (caudate, cortex, nucleus accumbens, hypothalamus, hippocampus) were also associated with AUDIT-C (Supplementary Tables 19). S-PrediXcan for AUDIT-C and AUDIT-P (Supplementary Table 20) revealed that lower predicted *RFC1* expression in the cerebellum and hemisphere, respectively, was associated with higher scores on both AUDIT-C (p = 7.84 × 10^−7^) and AUDIT-P (p = 1.54 × 10^−6^).

### Genetic correlations

We used LDSC to evaluate evidence for genetic correlations between our three primary traits (AUDIT total score, AUDIT-C, and AUDIT-P) and numerous other traits for which GWAS summary statistics were available; these included alcohol and substance use traits, personality and behavioral traits, psychiatric disorders, blood lipids, and brain structure volumes (**Supplementary Tables 3-5 and** Figure 2).

**Figure 2.**
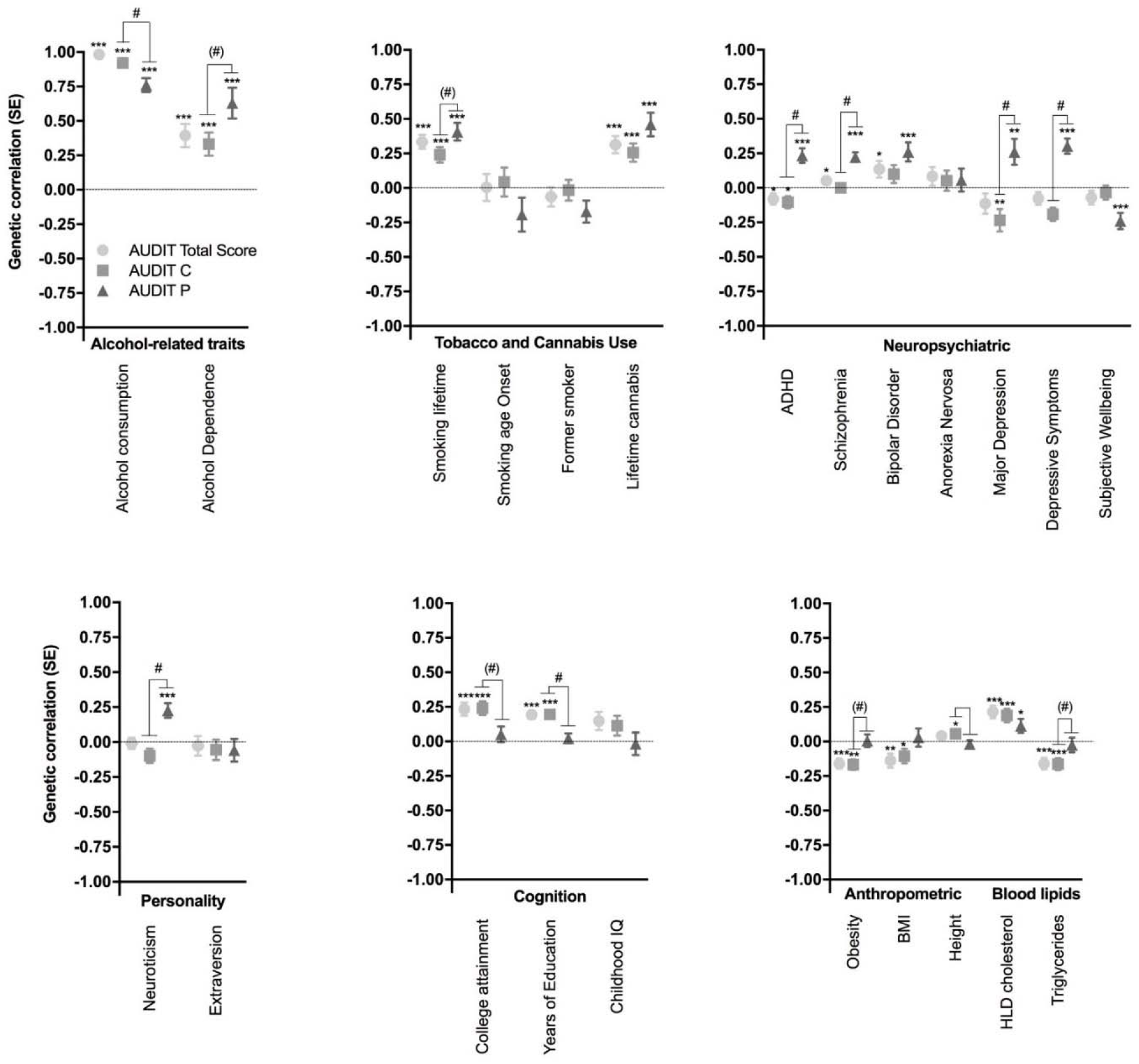
Genetic correlations between the three AUDIT phenotypes (total score, AUDIT-C, AUDIT-P) and several traits measured in independent cohorts as described in the Supplementary Tables 3-5: alcohol-related traits, tobacco and cannabis use, neuropsychiatric, personality, cognition, anthropomorphic and blood lipids. ADHD, attention-deficit/hyper-activity disorder; BMI, body mass index; SE, standard error; IQ, intelligence quotient; HDL, high-density lipoprotein. * p < 0.05, ** p < 0.01, *** p < 0.0001; # AUDIT-P vs AUDIT-C, p < 0.01 FDR 5%, (#) AUDIT-P vs AUDIT-C, p < 0.05

As expected, AUDIT-C and AUDIT-P were highly genetically correlated (r_g_ = 0.70, p = 4.1 × 10^−70^). AUDIT scores (AUDIT total score, AUDIT-C, and AUDIT-P) showed strong genetic correlations with alcohol consumption from two other studies (r_g_ = 0.76-0.96, p < 2.3 × 10^−9^). Many of the genetic correlations with AUDIT-P were significantly different from the correlations with AUDIT-C (**Supplementary Table 6**). The AUDIT-C had a significantly stronger (p = 8.02 × 10^−3^) correlation with alcohol consumption (r_g_ = 0.92, p = 7.0 × 10^−164^) than did AUDIT-P (r_g_ = 0.76, p = 2.7 × 10^−52^). In contrast, AUDIT total and AUDIT-C scores were only modestly correlated with alcohol dependence (r_g_ = 0.39 and 0.33 respectively, p < 8.2 × 10^−5^), whereas AUDIT-P showed a nominally stronger genetic correlation with alcohol dependence (r_g_ = 0.63, p = 1.8 × 10^−8^; AUDIT-P vs AUDIT-C, p = 0.033; see **Supplementary Table 6**).

We detected positive genetic correlations between AUDIT scores (AUDIT total, AUDIT-C, AUDIT-P) and other substance use phenotypes, including lifetime smoking (r_g_ = 0.24-0.41, p < 1.6 × 10^−5^) and cannabis use (r_g_ = 0.26-0.46, p < 1.1 × 10^−4^). We also observed a positive genetic correlation between AUDIT-P and cigarettes per day (r_g_ = 0.28, p = 4.0 × 10^−3^).

Several psychiatric disorders and related traits were positively genetically correlated with AUDIT-P scores, including schizophrenia (r_g_ = 0.22, p = 3.0 × 10^−10^), bipolar disorder (r_g_ = 0.26, p = 1.5 × 10^−4^), ADHD (r_g_ = 0.23, p = 1.1 × 10^−5^), and major depressive disorder (MDD, r_g_ = 0.26, p = 5.6 × 10^−3^). Intriguingly, AUDIT-C was negatively correlated with MDD (r_g_ = −0.23, p = 3.7 × 10^−3^) and ADHD (r_g_ = −0.10, p = 1.8 × 10^−2^), whereas AUDIT-P showed positive genetic correlations with these same disease traits (r_g_ = 0.26, p = 5.6 × 10^−3^; r_g_ = 0.24, p = 1.1 × 10^−5^).

We observed a positive genetic correlation between AUDIT-P scores and depressive symptoms (r_g_ = 0.30, p = 3.0 × 10^−8^) and neuroticism (r_g_ = 0.18, p = 2.6 × 10^−4^), and a negative genetic correlation with subjective well-being (r_g_ = −0.24, p = 4.0 × 10^−5^).

We observed positive genetic correlations between AUDIT total score, AUDIT-C and education, college attainment, and cognitive ability (r_g_ = 0.19-0.24, p < 1.5 × 10^−5^). The AUDIT-P genetic correlations with the same education and college attainment were near zero, and were significantly lower than AUDIT C and AUDIT total or education traits (**Supplementary Table 6**).

There were negative genetic correlations between AUDIT total and AUDIT-C scores and obesity (r_g_ = −0.16−0.17, p < 1.1 × 10^−5^), similar to previous reports regarding AUDIT total score (10) and alcohol consumption (9). In contrast, obesity was not significantly genetically correlated with AUDIT-P scores (r_g_ = 0.01, p = 0.90). Similarly, HDL cholesterol and triglycerides were genetically correlated with AUDIT total score and AUDIT-C (r_g_ = 0.19−22, p < 9.3 × 10^−5^, r_g_ = −0.16, p < 1.0 × 10^−4^ respectively), but this association was not found for AUDIT-P (r_g_ = 0.11, p = 2.2 × 10^−2^, rg = −0.03, p = 6.4 × 10^−1^). Obesity showed significantly different correlations with both AUDIT-P and AUDIT-C (**Supplementary Table 6**)

### Dichotomizing AUDIT total score to more closely approximate alcohol dependence

As AUDIT can be rapidly ascertained in large populations, we explored methods for dichotomizing AUDIT total score that optimized the genetic correlation with DSM-IV alcohol dependence (13). Higher genetic correlations with alcohol dependence were observed as the control threshold was increased from 2 to 4, and with increasingly stringent case cut-offs (Figure 3 and **Supplementary Table 2**). The highest genetic correlation was observed for cases with AUDIT total score ≥20 and controls ≤4 (r_g_ = 0.90, SE = 0.25, p = 3.0 × 10^−4^), however, this highly stringent threshold produced very few cases (N=1,290). The standard error of the estimate is much larger at more stringent case thresholds and therefore these estimates should be interpreted with caution. Defining cases as ≥12 yielded an r_g_ of 0.82 (SE = 0.18, p = 3.2 × 10^−6^) while retaining more than 7 times as many cases (N=9,130), these genetic correlations were not significantly different from those obtained using cases ≥20 and controls ≤4 (p = 0.80).

**Figure 3.**
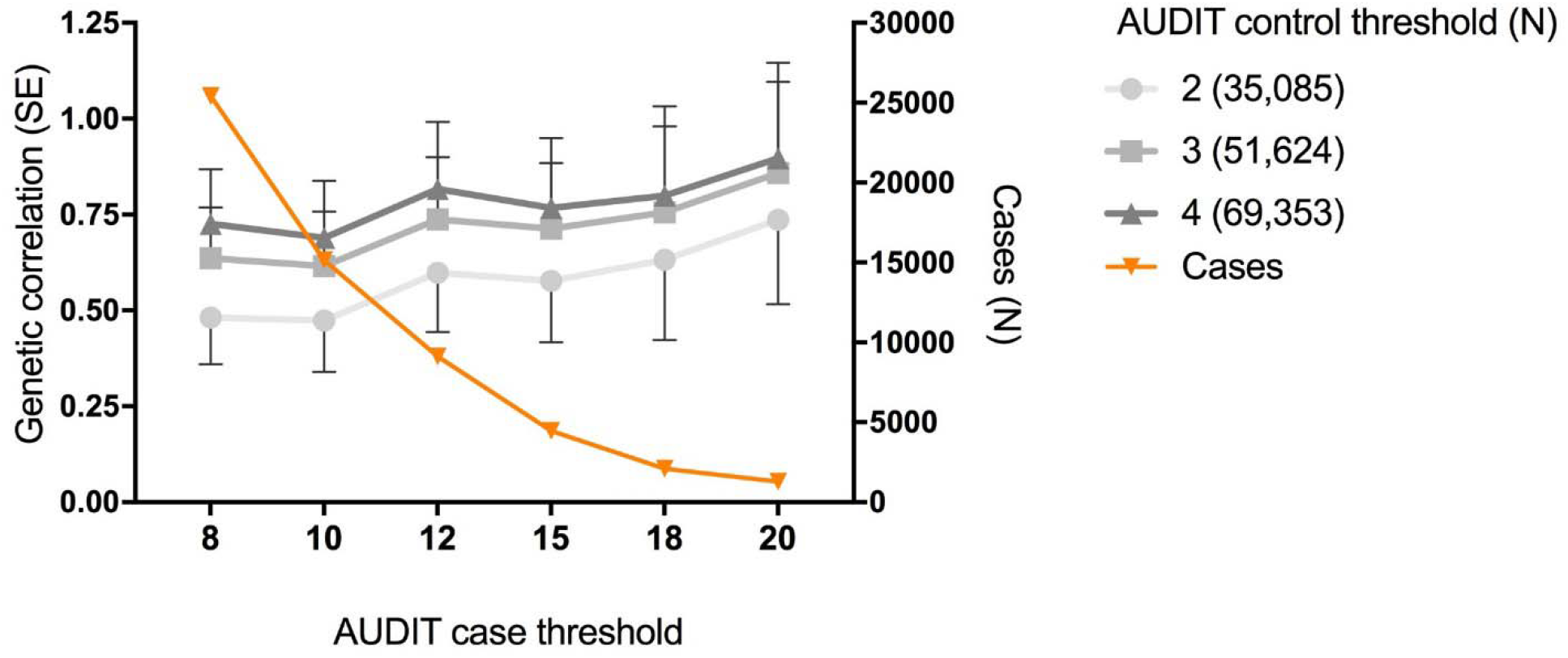
Genetic correlations between AUDIT cases (8 [N = 25,423], 10 [N = 15,151], 12 [N = 9,130], 15 [N = 4,471], 18 [N = 2,099], 20 [N = 1,290]) vs controls (2, 3, 4) in the UK Biobank and DSM-IV derived alcohol dependence from the Psychiatric Genetics Consortium. The orange line is a visualization of the number of cases used at each threshold, corresponding to the N on the right hand y-axis.

## Discussion

We have presented the largest GWAS meta-analysis of AUDIT total score to date, using large population-based cohorts from UKB and 23andMe. We identified novel associations with AUDIT total score; the genes located in these regions include *JCAD* and *SLC39A8*. We found evidence for association in several loci previously associated with alcohol use via single-variant and gene-based analyses (i.e. *KLB, GCKR, CADM2*). The SNP heritability of all AUDIT phenotypes ranged from 9-12% demonstrating that common genetic factors account for a prominent proportion of the variation in alcohol use phenotypes. Furthermore, we showed that there is shared genetic architecture between AUDIT scores and other alcohol and substance use phenotypes. AUDIT-P showed a positive genetic correlation with several psychiatric diseases, distinguishing AUDIT-P from AUDIT-C. Finally, using LDSC, we identified thresholds for dichotomizing AUDIT total score (AUDIT score ≥12 to define cases, and ≤4 to define controls) that maximize the genetic correlation with alcohol dependence while retaining a large number of participants.

Our top GWAS hits replicated previous association signals for alcohol use traits. The strongest associations with AUDIT score in this study spanned the alcohol metabolism genes on chromosome 4q23 (10). Variants in this region were associated with AUDIT total score, AUDIT-C and AUDIT-P, demonstrating that alcohol metabolism is a risk factor for both alcohol consumption and problematic use. The second strongest signal, also associated with the three AUDIT phenotypes, is located in *KLB*, confirming the robust association of this gene with both alcohol consumption (9, 11, 12) in humans, and in mice (12). However, the biology of this locus could be more complex than previously described. Although the credible set analysis suggested that the more probable causal variants are all located on the first intron of *KLB*, one of these variants, rs11940694, is an eQTL for *RFC1* expression in the brain, and S-PrediXcan analysis predicted that lower expression of *RFC1* in the brain is associated with higher predicted AUDIT (AUDIT-C and AUDIT-P) scores. Interestingly, a gene in the complex GWAS signal on chromosome 19, Fibroblast growth factor 21 (*FGF21*), was associated with AUDIT (AUDIT total score, AUDIT-C, AUDIT-P) at the gene-based level (**Supplementary Table 17**). *FGF21* regulates sweet and alcohol preference in mice as part of a receptor complex with β-Klotho (*KLB*) in the central nervous system (33). Additionally, we replicated the association between rs1260326 in the gene *GCKR* and alcohol consumption (9, 11), here associated with AUDIT total score and AUDIT-C. Other loci previously associated with alcohol consumption include *CADM2* (9), which was associated at the gene-based level for all three AUDIT traits. Here, the burden analysis suggests that multiple (rare and common) variants are necessary to explain the association signal. Intriguingly, several of the novel associations with AUDIT scores were mapped to highly pleiotropic genes (*MAPT, FUT2, SLC39A8*)(27).

Genetic analysis of the AUDIT subsets revealed evidence of distinct genetic architecture between AUDIT-C and AUDIT-P (alcohol consumption vs. problem use), with support from the gene-based (**Supplementary Figures 16** and **17**), S-PrediXcan (**Supplementary Tables 19** and **20**), and genetic correlation analyses (Figure 2). Furthermore, AUDIT-P showed a strong genetic correlation with alcohol dependence (13). In contrast, AUDIT-C had a stronger genetic correlation with alcohol consumption. Thus, partitioning AUDIT scores into different subsets (alcohol consumption vs problem use) may disentangle genetic factors that contribute to different aspects of AUD vulnerability.

Genetic overlap was observed for all measures of AUDIT and other substance use traits, including lifetime tobacco and cannabis use, as we previously reported (10, 36, 37), demonstrating that genetic risk factors for high AUDIT scores overlap with increased consumption of multiple drug types.

We found several significant differences between the genetic correlations with AUDIT-P and AUDIT-C. These differences were particularly pronounced for psychiatric and behavioral traits. AUDIT-P was positively genetically correlated with psychopathology (schizophrenia, bipolar disorder, MDD, ADHD), personality traits including neuroticism, and regional brain volumes. These associations have previously been observed at the phenotypic level; AUDs commonly co-occur in individuals with schizophrenia (38), bipolar disorder (39), MDD (40) and adult ADHD (41). Intriguingly, genetic risk for high AUDIT-C score was *negatively* correlated with MDD and ADHD demonstrating that a distinct genetic component of AUDIT-P is shared with genetic risk for psychiatric disease. Regional volume abnormalities in subcortical brain regions of AUD individuals have been reported (42–44), however, it is unclear whether these alterations are a result of high alcohol drinking or a pre-existing susceptibility. We identified a positive genetic correlation between AUDIT-P and increased caudate volume; however, the majority of studies report reductions in regional brain volumes associated with AUD.

For AUDIT total score and AUDIT-C we showed positive genetic correlations with educational attainment and cognitive ability and negative genetic correlations with obesity, consistent with earlier reports (9, 10). These associations were not observed for AUDIT-P. Similarly, HDL cholesterol showed a significant positive correlation, and triglycerides a negative correlation, with AUDIT total score and AUDIT-C, but not AUDIT-P. These patterns were previously observed for alcohol consumption (9). We could speculate that these differences may be linked to socioeconomic status (SES). Alcohol consumption is often higher in individuals with higher SES (45), whereas alcohol-related problems, such as binge drinking (46) and alcohol related mortality (47), are more prevalent in individuals with lower SES. Furthermore, individuals with low SES are more likely to have AUDs with psychiatric co-morbidities (48). Consistent with this idea, we find positive genetic correlations between AUDIT-C and education, a trait correlated with SES (49), and positive genetic correlations between AUDIT-P and psychopathology. Our findings provide further evidence that different dimensions of alcohol use associate differently with behavior and that these differences may have a biological underpinning.

A clinical diagnosis of AUD is often required to define cases for genetic studies. An alternative strategy would be to use AUDIT to infer AUD case status; however, it has not been clear whether and how to perform meta-analyses between AUDIT scores and alcohol dependence. A GWAS meta-analysis for AUDIT and alcohol dependence could be simplified if a threshold could be used to define cases and controls based on AUDIT scores, an approach that was used by Mbarek et al. (50). We have provided empirical evidence about genetic correlations between AUDIT and alcohol dependence using dichotomized AUDIT scores, and found thresholds for AUDIT that produced high genetic correlations with AUD (Figure 3). Genetic correlations increased as the upper threshold for cases was made more stringent, although the standard errors for all of these estimates were overlapping. The genetic correlation with alcohol dependence appeared to asymptote when case status was defined as ≥12; therefore, this threshold could be used to define case status. We also considered various thresholds for defining controls and found that ≤4 produced a high genetic correlation with alcohol dependence while also retaining the largest number of subjects.

Our study is not without limitations. AUDIT specifically asks about the past year, and thus may not capture information on lifetime alcohol use and misuse. This is suboptimal for genetic studies because it effectively measures a recent state rather than a stable trait. Measures capturing drinking and AUD across the lifespan may be preferable. Also, although mean scores for the AUDIT-C dimension were 4.24, the mean of the AUDIT-P dimension was considerably lower (0.75). Thus, we were not able to perform a more refined categorization (e.g. 3 subsets: consumption [items 1-3], dependence [items 4-6], hazardous use [items 7-10]) as fewer individuals endorsed the items comprising AUDIT-P (see **Supplementary Table 7**).

Furthermore, our study uses data from UKB and 23andMe research participants, who were volunteers not ascertained for AUD, and hence our findings may not generalize to other populations showing higher rates of alcohol use and dependence. Additional alcohol-related phenotypes (e.g. age at first use; patterns of alcohol drinking, including binge drinking) could be used in subsequent genetic studies to identify additional sources of genetic vulnerability for AUD. Lastly, we offered guidelines to identify cases to use in genetic studies of AUD (i.e. AUDIT score ≥12) based on genetic correlations; however, these recommendations are not intended to determine thresholds for diagnosing dependence in a clinical setting. Future studies will be able to test whether using AUDIT as a surrogate for AUD will be beneficial for gene discovery. In addition, several studies have argued that lower thresholds should be used for females, which has not been addressed in the present study.

We have reported the largest GWAS of AUDIT ever undertaken. We replicated previously identified signals (i.e. *ADH1B, ADH1C; KLB; GCKR*), and identified novel GWAS signal (i.e. *JACD, SLC39A8*) associated with AUDIT. We show that different portions of the AUDIT (AUDIT-C, AUDIT-P) correlate with distinct traits, which will aid in dissecting genetic vulnerability towards alcohol use and dependence. The genetic factors that predispose to high alcohol consumption inevitably overlap with those for problem drinking, as heavy drinking is generally a prerequisite for the development of hazardous use and dependence. However, not everyone who consumes alcohol experiences the same level of harmful consequences. By studying the different subsets of AUDIT, we identify genetic factors that may be specific to problem drinking. Larger studies of cohorts with a wider range of AUDIT-P scores are required to both replicate and expand these findings. Finally, we describe an alternative strategy to rigorous ascertainment for genetic studies of AUD, i.e. AUDIT score ≥12 to define cases and ≤4 to define controls, which could be used to achieve large sample sizes in a cost-efficient manner.

### URLs

UK Biobank: http://www.ukbiobank.ac.uk/

FUMA: http://fuma.ctglab.nl/

LD score software: https://github.com/bulik/ldsc/

LDHub: http://ldsc.broadinstitute.org/

METAL: https://genome.sph.umich.edu/wiki/METAL

S-PrediXcan: https://github.com/hakyimlab/S-PrediXcan-Working

The NHGRI GWAS Catalog: http://www.genome.gov/gwastudies/

Regulome DB database: http://www.regulomedb.org/

PredictDB Data Repository: http://predictdb.hakyimlab.org/

Credible set estimation method, R script: https://github.com/hailianghuang/FM-summary

## Data availability

We have provided summary statistics for the top 10,000 SNPs (Supplementary Data Set). Full GWAS summary statistics for the 23andMe dataset will be made available through 23andMe to qualified researchers under an agreement with 23andMe that protects the privacy of the 23andMe participants. Interested investigators should for more information. GWAS summary statistics for the UKB GWAS of AUDIT scores will be available on request.

## Acknowledgements

We would like to thank the research participants and employees of 23andMe for making this work possible. S.S-R was supported by the Frontiers of Innovation Scholars Program (FISP; #3-P3029), the Interdisciplinary Research Fellowship in NeuroAIDS (IRFN; MH081482) and a pilot award from DA037844. This research has been conducted using the UK Biobank Resource: application number 4844 and was supported by a Wellcome Trust Strategic Award ‘Stratifying Resilience and Depression Longitudinally’ (STRADL) (Reference 104036/Z/14/Z), and by the Medical Research Council- and Biotechnology and Biological Sciences Research Council-funded Centre for Cognitive Ageing and Cognitive Epidemiology (Reference MR/K026992/1).

The authors report no conflict of interest

